# A native phosphoglycolate salvage pathway of the synthetic autotrophic yeast *Komagataella phaffii*

**DOI:** 10.1101/2023.09.30.560291

**Authors:** Michael Baumschabl, Bernd M. Mitic, Christina Troyer, Stephan Hann, Özge Ata, Diethard Mattanovich

## Abstract

Synthetic autotrophs can serve as chassis strains for bioproduction from CO_2_ as a feedstock to take measures against the climate crisis. Integration of the Calvin–Benson–Bassham (CBB) cycle into the methylotrophic yeast *Komagataella phaffii* (*Pichia pastoris*) enabled it to use CO_2_ as the sole carbon source. The key enzyme in this cycle is ribulose-1,5-bisphosphate carboxylase/oxygenase (RuBisCO) catalyzing the carboxylation step. However, this enzyme is error prone to perform an oxygenation reaction leading to the production of toxic 2-phosphoglycolate. Native autotrophs have evolved different recycling pathways for 2-phosphoglycolate. However, for synthetic autotrophs, no information is available for the existence of such pathways. Deletion of *CYB2* in the autotrophic *K. phaffii* strain led to the accumulation of glycolate, an intermediate in phosphoglycolate salvage pathways, suggesting that such a pathway is enabled by native *K. phaffii* enzymes. ^13^C tracer analysis with labeled glycolate indicated that the yeast pathway recycling phosphoglycolate is similar to the plant salvage pathway. This orthogonal yeast pathway may serve as a sensor for RuBisCO oxygenation, and as an engineering target to boost autotrophic growth rates in *K. phaffii*.

## Introduction

The Calvin–Benson–Bassham (CBB) cycle is responsible for the vast majority of carbon fixed on our planet. The key enzyme and rate limiting step of this cycle is ribulose-1,5-bisphosphate carboxylase/oxygenase (RuBisCO), which catalyzes the carbon dioxide fixation to ribulose-1,5-bisphosphate. Every year around 10^14^ kg carbon dioxide is fixed by RuBisCO (Hayer-Hartl and Hartl, 2020). With an average turn over number of around 3 s^-1^ RuBisCO is a very slow enzyme. Plants therefore produce huge amounts of this protein, with a RuBisCO content of around 30% in C4 plants and up to 50% of soluble protein in C3 plants (Feller et al., 2008). There are estimates that the total mass of RuBisCO proteins on our planet is one gigaton (Bar-On and Milo, 2019). These huge amounts of protein are necessary because it easily gets inhibited by a range of other sugar phosphates. These inhibitors can be released by the action of specific RuBisCO activases (Hauser et al., 2015; Parry et al., 2008).

Another aspect which reduces the efficiency of the RuBisCO protein is the fact that it also reacts with oxygen instead of CO_2_ which reduces the carbon fixation rate. The tendency to perform this unfavorable oxygenation reaction varies between different types of RuBisCO. In general, the RuBisCO protein family can be subdivided into 4 types where type I and II are the predominant forms in organisms performing the CBB cycle. Type I proteins consisting of 8 large and 8 small subunits are present in plants, cyanobacteria and proteobacteria while type II proteins form homodimers of a large subunit only and are mainly found in proteobacteria (Hauser et al., 2015; Tabita et al., 2008). Type II proteins have often higher turnover numbers compared to type I proteins but lack their higher specificity to CO_2_. Overall there is an inverse correlation between the specificity and the turnover rate (Tcherkez et al., 2006).

Besides adapting the protein itself, organisms developed carbon concentrating mechanisms to increase the concentration of CO_2_ in the proximity of RuBisCO to reduce the oxygenation rate (Raven et al., 2008). C4 plants can store CO_2_ by carboxylation of phosphoenolpyruvate (PEP) to oxaloacetate which is later converted to malate and afterwards transported to bundle sheet cells and decarboxylated close to the RuBisCO protein to increase the local concentration of CO_2_ (**Fig. 1 A**) (Keeley and Rundel, 2003). Cyanobacteria and some chemoautotrophic bacteria developed protein based microcompartments called carboxysomes which encapsulate RuBisCO. Organisms carrying carboxysomes import bicarbonate using transmembrane pumps and this bicarbonate is then converted back to CO_2_ in the carboxysomes using carbonic anhydrase (Cameron et al., 2013; Yeates et al., 2008). Bicarbonate is the dominant form of dissolved carbon species in aquatic environments making it the predominant molecule to be imported by the cells as a carbon source. This system is very effective and can increase the CO_2_ concentration up to 1000-fold (**Fig. 1 B**) (Wang et al., 2015).

**Figure 1:**
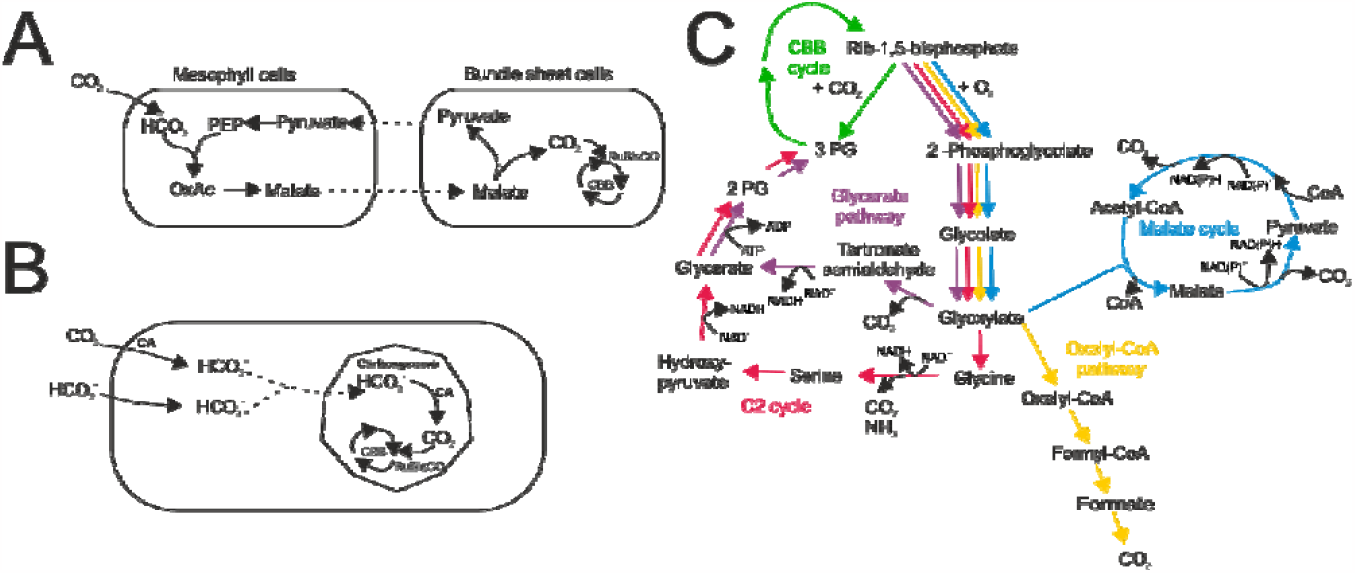
Phosphoglycolate salvage pathways and carbon concentrating mechanisms. (A) Carbon concentrating mechanism of C4 plants. (B) Carbon concentrating mechanism of organisms containing carboxysomes. (C) Overview of different phosphoglycolate salvage pathways: C2 cycle in red, glycerate pathway in purple, oxalyl CoA pathway in yellow and the malate cycle in blue. CBB: Calvin– Benson–Bassham cycle, 2-PG: 2-phosphoglycerate, 3-PG: 3-phosphoglycerate, Rib-1,5-bisphosphate: ribulose-1,5-bisphosphate, CoA: coenzyme A, PEP: phosphoenolpyruvate, OxAc: oxaloacetate, CA: carbonic anhydrase

Even though various organisms harboring the CBB cycle perform carbon concentrating mechanisms to improve their carbon capture efficiency, the oxygenation reaction is always present to some degree. Oxygenation of ribulose 1,5-bisphosphate produces one molecule 3-phosphoglycerate (3-PG) and one molecule 2-phosphoglycolate. 2-phosphoglycolate is reported to be toxic because it can inhibit at least two key enzymes of the central carbon metabolism, triose phosphate isomerase and phosphofructokinase (Flügel et al., 2017; Hall et al., 1987). Therefore, organisms came up with pathways recycling this accumulating by-product. Among different types of organisms different pathways for recycling have evolved. In plants, most algae and cyanobacteria recycling is performed by the so called C2 cycle (**Fig. 1 C** red line). In this cycle 2-phosphoglycolate is dephosphorylated in the chloroplasts, then oxidized by glycolate oxidase to glyoxylate. The next step, transamination of glyoxylate to glycine, can be performed by two parallel reactions with different molecules used as amino group donors: serine:glyoxylate aminotransferase (SGT) and glutamate:glyoxylate aminotransferase (GGT). In the mitochondria two molecules glycine are converted to serine which releases ammonia and CO_2_. The produced serine is then transported back to the peroxisomes where it donates its amino group again to a glyoxylate molecule catalyzed by SGT. The deamination product hydroxypyruvate is then transported to the cytosol and reduced to glycerate. The last step in this cycle is the phosphorylation to 3-phosphoglycerate performed in the chloroplast which enables the re-entry into the CBB cycle (Bauwe et al., 2010). Taking the required energy for the refixation of the deaminated amino group into account the photorespiration pathway in plants consumes 3.5 ATP and 2 NADH equivalents and releases 0.5 molecules of CO_2_ per RuBisCO oxygenation (Walker et al., 2016).

In cyanobacteria two different pathways responsible for the salvage of 2-phosphoglycolate have been identified besides the C2 cycle. In one of these pathways, the glycerate pathway, 2 molecules of glyoxylate are converted to tartronate semialdehyde, further reduced to glycerate and phosphorylated ending up in 3-PG avoiding the release of ammonia compared to the C2 cycle (**Fig. 1 C** violet line). The other pathway catalyzes the complete oxidation of 2-phosphoglycolate to CO_2_ via oxalate and formate in combination with harvesting of energy in the form of NADH (**Fig. 1 C** yellow line) (Eisenhut et al., 2008, 2006). In the chemolithotrophic bacterium *Cupriavidus necator* the glycerate pathway is the main contributor to the recycling of 2-phosphoglycolate but another pathway was identified, the malate cycle which fully oxidizes glycolate to CO2 (**Fig1. C** blue line) (Claassens et al., 2020) while harvesting energy in the form of 2 molecules NADH per glycolate oxidation.

All organisms which naturally perform the CBB cycle perform at least one of the possible routes for recycling of 2-phosphoglycolate. Lately the CBB cycle was successfully integrated in the two model organisms *Escherichia coli* (Gleizer et al., 2019) and *Komagataella phaffii* (Gassler et al., 2020). In both approaches no specific pathway for the phosphoglycolate recycling was integrated. Cells were able to grow efficiently with CO_2_ as sole carbon source leading to the assumption that the strains came up with a phosphoglycolate salvage pathway based on native enzymes. The aim of this study was to identify and characterize the phosphoglycolate salvage in the synthetic autotrophic *K. phaffii* strain.

## Results

### Are synthetic autotrophs sensitive to oxygen?

We recently created a synthetic autotrophic *K. phaffii* strain by the deletion of DAS1 and DAS2 which blocks the assimilation of methanol using the xylulose-5-phosphate cycle as well as by the integration of six genes of the CBB cycle including the RuBisCO protein CbbM of *Thiobaccillus denitrificans* and Prk from *Spinacia oleracea* (Gassler et al., 2020). This strain was able to grow on CO_2_ as sole carbon source and methanol as energy source. The key enzyme of the CBB cycle is RuBisCO fixing the carbon dioxide to ribulose-1,5-bisphosphate, but also catalyzing the reaction with oxygen. Many organisms performing the CBB cycle have come up with carbon concentrating mechanisms to locally increase CO_2_ concentration to reduce the rate of oxygenation. These mechanisms are lacking in our synthetic autotrophic yeast. Therefore, we wanted to evaluate the effect of the oxygen concentration on growth rate of this strain. Bioreactor cultivations with oxygen concentrations in the inlet gas between 2.5% and 20% were performed and growth and dissolved oxygen were monitored.

We clearly saw a difference between the different oxygen concentrations in the inlet gas (**Fig 2A**). The two lower oxygen concentrations of 5% and 10% resulted in faster growth compared to the higher oxygen levels in the inlet gas. The lowest oxygen concentration of 2.5% resulted in slightly lower growth compared to 5 and 10%. A similar picture was observed in an engineered version (Gassler et al., 2022) of this strain which enables it to reach faster autotrophic growth. Here the only difference in terms of oxygen sensitivity was a slightly increased oxygen demand of this strain (**Supp. Fig 1A**), with an optimum oxygen concentration around 15%.

**Figure 2:**
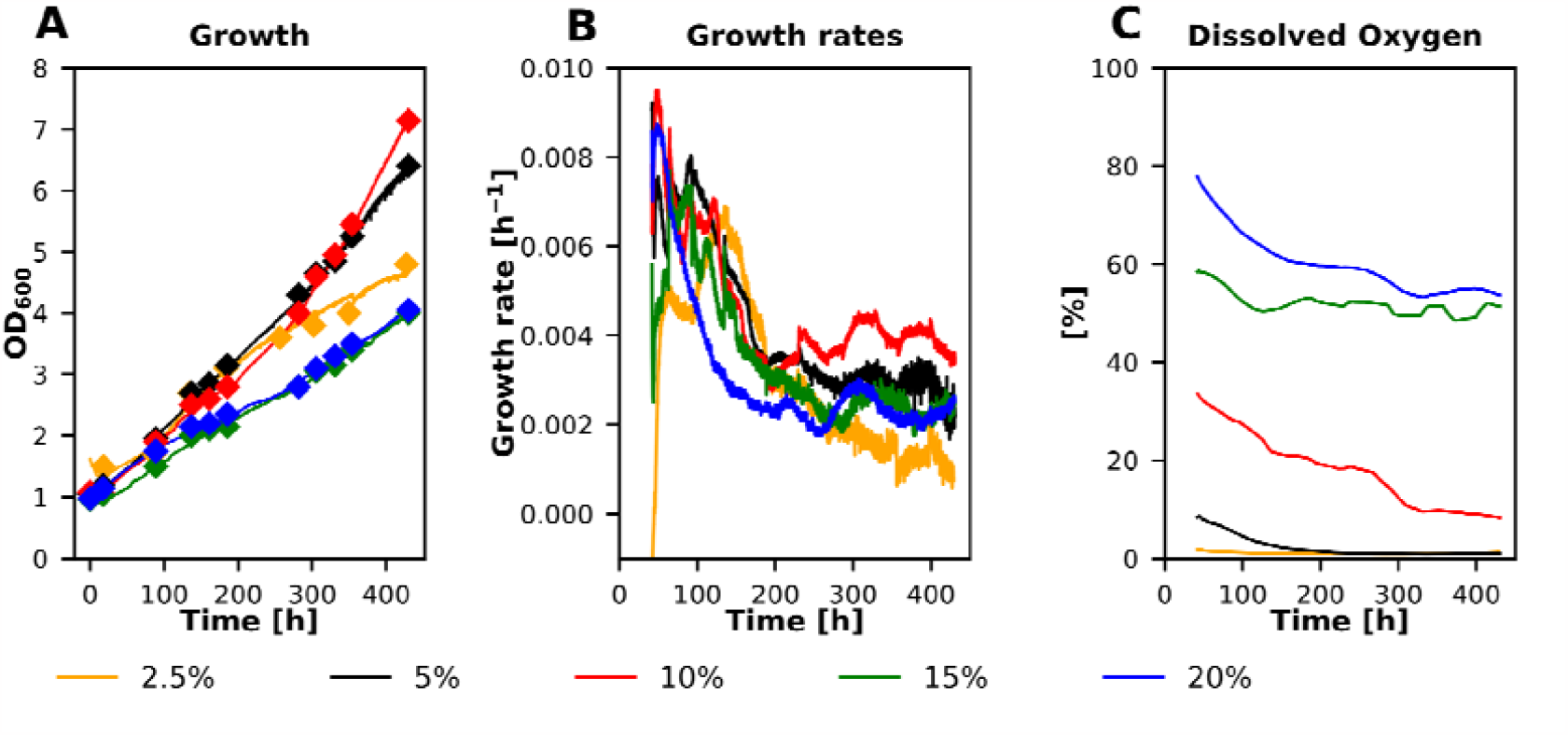
Bioreactor cultivations using different oxygen levels in the inlet air to test their influence on growth. (A) Diamonds: offline OD_600_ measurements and solid line online OD probe to monitor growth, (B) calculated growth rates and (C) dissolved oxygen concentrations of 5 different fermentations using oxygen concentrations in the inlet air from 2.5% to 20% v/v. Cultivations were performed at 30°C with a constant stirrer speed of 300 rpm and a CO_2_ concentration in the inlet air of 5%.

### A single gene deletion blocks phosphoglycolate salvage

While reducing the oxygen concentration improved the growth rate of the synthetic autotroph yeast, it can also grow efficiently at ambient oxygen levels. This suggests that this synthetic autotroph is able to perform all reactions necessary to recycle 2-phosphoglycolate. When we engineered lactic acid production into this strain in a previous study, we deleted *CYB2* encoding for an L-lactate cytochrome-c oxidoreductase which reduces lactate to pyruvate in the mitochondrial intermembrane space. Besides decreasing lactic acid reassimilation as intended, this knockout obviously interrupted the 2-phosphoglycolate salvage pathway, as the deletion strain secreted glycolate into the medium under autotrophic conditions (Baumschabl et al., 2022). This gave already a first hint that there is a pathway responsible for the recycling of the formed phosphoglycolate and that the oxidoreductase Cyb2 is involved in this.

To identify the possible route of 2-phosphoglycolate salvage formed by the oxygenation reaction of RuBisCO we perfomed ^13^C tracer experiments with fully ^13^C labeled glycolate. To make sure that glycolate can be used as tracer metabolite, the synthetic autotrophic yeast strain and variants of this strain overexpressing *CYB2* under the control of a medium or weak promoter were cultivated using glycolate as the only carbon source. None of the tested strains were able to grow on glycolate (**Fig. 3 A**), but all strains could assimilate it. When *CYB2* was overexpressed, a higher assimilation rate was observed (**Fig. 3B**). These results confirmed that glycolate is assimilated in the metabolism and ^13^C from glycolate will therefore be incorporated.

**Figure 3:**
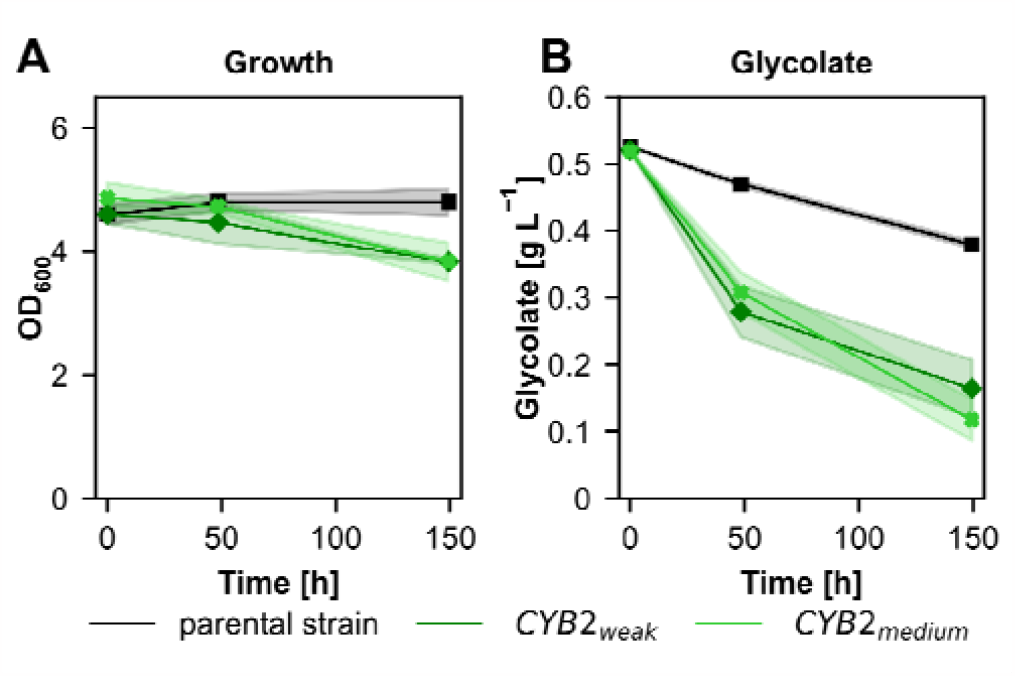
Cultivations using glycolate as sole carbon source. (A) Growth and (B) glycolate concentrations in the supernatant of the synthetic autotrophic *K. phaffii* strain (parental strain) and *CYB2* overexpression strains using weak or medium strength promoters. Cultivations were performed at 30°C and ambient CO_2_ concentrations in the atmosphere. Solid lines indicate the mean and shades the standard deviation of 3 biological replicates.

### Identification of the native phosphoglycolate salvage pathway

Glycolate can be efficiently assimilated by the autotrophic *K. phaffii* strain which is a prerequisite for its use as a tracer metabolite to further investigate the route of phosphoglycolate salvage. Therefore, three different strains, the parental autotrophic strain, a *CYB2* knockout strain and the *CYB2* overexpression strain using the medium strength promoter, were cultivated on fully labeled ^13^C glycolate for 48 hours. Samples were taken after 3, 24 and 48 hours for the determination of the isotopologue distribution of various intracellular metabolites using GC-TOFMS. If a metabolite was involved in the phosphoglycolate salvage pathway, it would show a decrease in the “M 0” isotopologue (^12^C only) (blue bars in Fig 4) and an increase in the higher mass isotopologues, showing that ^13^C from labeled glycolate is incorporated into the respective metabolite. Metabolites with the highest ^13^C content are supposed to be located at the start of a pathway. When interpreting the labeling data, it had to be considered that all strains produce ^12^C glycolate via oxygenation of ribulose 1,5-bisphosphate catalyzed by the expressed RuBisCO, which can reduce the incorporation of ^13^C atoms.

**Figure 4:**
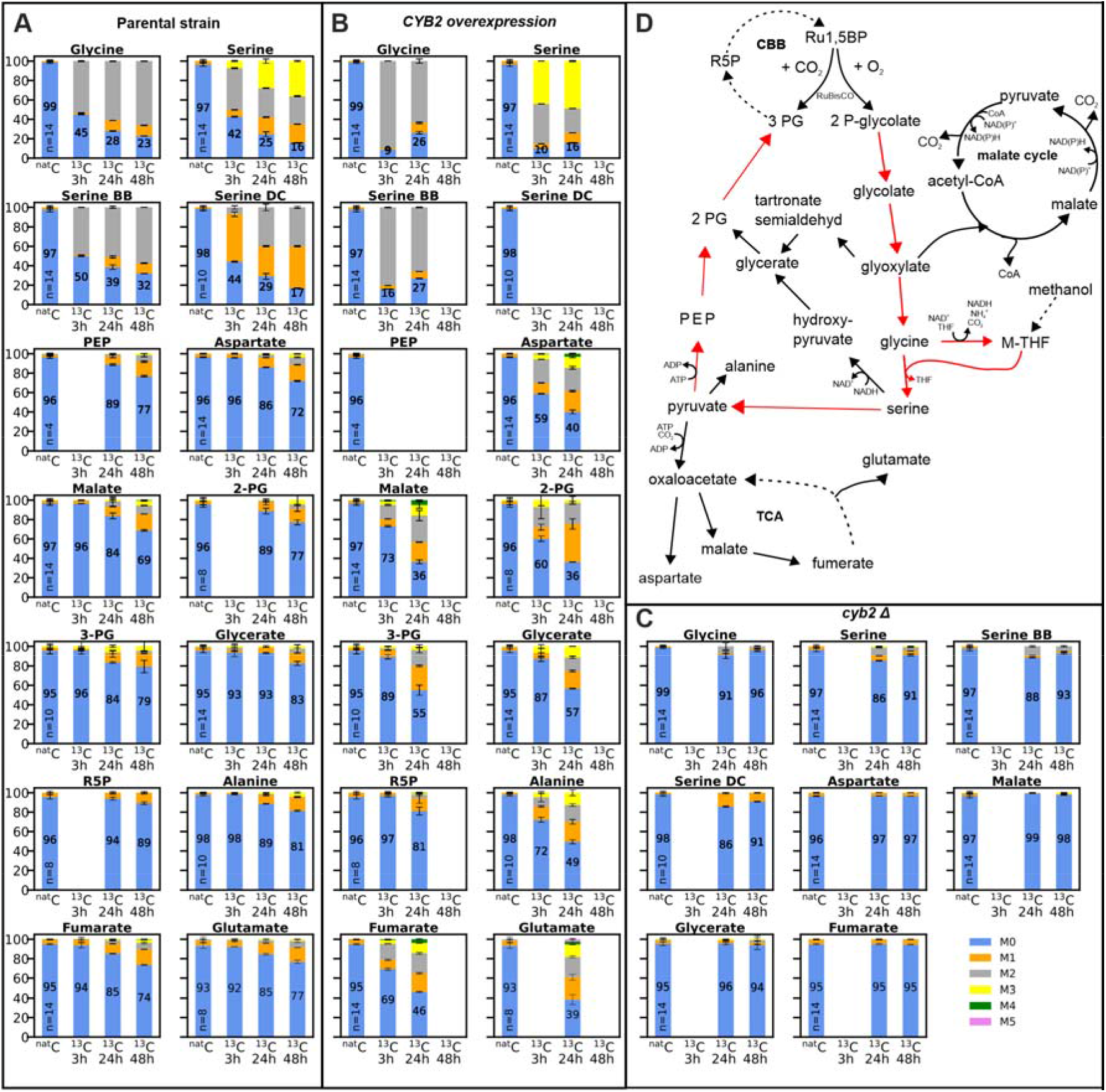
Carbon isotopologue distribution of intracellular metabolites resulting from tracer experiments using fully labeled ^13^C glycolate as carbon source to reveal the phosphoglycolate salvage pathway in the synthetic autotroph K. phaffii strain. Results of (A) parental strain, (B) CYB2_medium_ overexpression strain and (C) CYB2 deletion strain. M0 to M5 corresponds to the number of ^13^C atoms present in the metabolite. The numbers in the blue bars indicate the isotopologue fraction of the M0 isotopologue of the respective sample. The number of biological replicates of the ^nat^C samples is indicated by ‘n=’ in the bars; 2 biological replicates were performed for all ^13^C labeled samples. Error bars represent the standard deviation. Serine BB denotes the amino acids backbone fragment including only the C1 and C2 carbon atom of serine. Serine DC denotes the decarboxylated molecule with the C1 carbon atom being cleaved off. For the parental strain all three, for the overexpression strain the first two and for the deletion strain the last two time points were measured. Data of the missing bars could not be evaluated as they did not match the quality criteria. Abbreviations: PEP: phosphoenolpyruvate, 2-PG: 2-phosphoglycerate, 3-PG: 3-phosphoglycerate, R5P: ribose 5-phosphate, Ru1,5BP: ribulose 1,5-bisphosphate, CBB: Calvin–Benson–Bassham cycle, M-THF: methylene tetrahydrofolate, TCA: tricarboxylic acid cycle, CoA: coenzyme A

In the parental strain pronounced incorporation of ^13^C atoms into serine and glycine was observed already after 3 hours (**Fig. 4A**), indicating that most of the glycolate is recycled via glycine and not oxidized in the malate cycle. The ^13^C content of both metabolites increased until the end of cultivation. Neither pyruvate nor hydroxypyruvate could be analyzed with the applied GC-TOFMS method, hence the metabolic step following serine had to be deduced from labeling patterns of adjacent metabolites. Significant incorporation of ^13^C into PEP, 2-PG, 3-PG and glycerate was observed but to a lower extent compared to serine and glycine. Ribose 5-phosphate showed only a small fraction of ^13^C labeled isotopologues. Metabolites related to the TCA cycle such as malate and fumarate showed a similar labeling degree as the phosphorylated hydroxycarboxylic acids. Since malate and aspartate showed similar labeling patterns, the labeling pattern of both metabolites probably originates from oxaloacetate and not from the malate cycle. Two additional fragments of serine were evaluated: the backbone fragment (BB) corresponding to the amino acid backbone of serine containing the C1 and C2 carbon atom only and the decarboxylated fragment (DC) which corresponds to the decarboxylated serine molecule containing the C2 and C3 carbon atom. The two fragments showed significantly different labeling patterns. BB had a much higher fraction of the fully labeled isotopologue M2 compared to DC which showed a high fraction of M1, i.e. one labeled carbon atom. This finding indicates that especially at the beginning of the cultivation the methylene group of M-THF, which is used for the synthesis of serine from glycine, had a high ^12^C content. At later stages of the cultivation, the fraction of the M2 isotopologue of the DC fragment increased indicating that labeled glycine is used for M-THF synthesis.

In the strain overexpressing *CYB2* the labeling pattern of the metabolites was similar but in general a higher incorporation of ^13^C into the metabolites was observed (**Fig. 4 B**). Already after 3 hours of cultivation on ^13^C glycolate glycine and serine were nearly fully labeled. The ^13^C content was reduced at the next sampling time point (24 h), because nearly all ^13^C glycolate was consumed and more ^12^C glycolate produced from RuBisCO was present (**Supplementary Fig. 2**). In 2-PG a higher incorporation of ^13^C was observed compared to 3-PG as well as glycerate, which showed similar labeling patterns. Again ribose 5-phosphate, the CBB intermediate evaluated here, showed only a small fraction of ^13^C labeled isotopologues. In addition, high ^13^C contents can be found in the TCA metabolites malate and fumarate as well as in aspartate and glutamate, obviously deriving from oxaloacetate.

In contrast to the other two strains, the strain harboring the deletion of *CYB2* resulted in a minor incorporation of ^13^C into glycine and serine (**Fig. 4 C**). All other analyzed metabolites did not show any significant incorporation of ^13^C labeled carbon. As glycine and serine are weakly labeled in this strain, we assumed an alternative gene to *CYB2* catalyzing the same reaction to be present. One candidate gene was *DLD1* which shows similarities to *C. necator* glycerate dehydrogenase being responsible for the oxidation of glycolate to glyoxylate. However, an overexpression of this gene did not result in any significant glycolate consumption (**Supplementary Fig 3**).

In summary, the tracer experiments confirmed that *CYB2* plays a major role in the recycling of 2-phosphoglycolate produced by the oxygenation reaction of RuBisCO. While the deletion of *CYB2* blocked the incorporation of ^13^C from glycolate almost completely, its overexpression boosted the flux of this pathway. Cyb2 is annotated as an L-(+)-lactate-cytochrome c oxidoreductase. Given the structural similarity of L-lactate and glycolate, the results can be interpreted such that Cyb2 also oxidizes glycolate to glyoxylate, taking over the role of glycolate oxidase and transferring the electrons to cytochrome *c* (Cunane et al., 2002). To evaluate if the natural activity of Cyb2 could lead to a bottleneck in phosphoglycolate recycling and therefore may limit the autotrophic growth rates, we cultivated *CYB2* overexpression strains using a medium and weak promoter under autotrophic conditions. We included also a cytosolic version of this protein to test if cytosolic location helps to boost the growth rate.

None of the tested constructs could increase growth under autotrophic conditions (**Fig. 5**). It seemed that the cytosolic version of *CYB2* with the mitochondrial signal sequence being deleted induced more stress and reduced growth rate even more.

**Figure 5:**
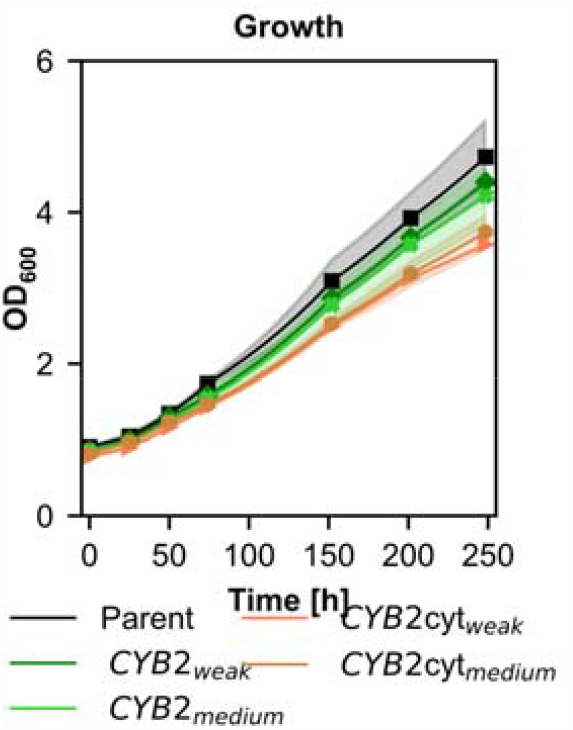
Autotrophic growth of the CYB2 overexpression strains compared to their parental strain. Cultivations were performed at 30°C and 5% CO_2_ concentrations in the atmosphere. Solid lines indicate the means, shades the standard deviation of the biological replicates.

### The *CYB2* deletion strain can serve as RuBisCO oxygenation test platform

Deletion of *CYB2* abolished the conversion of glycolate into glycine and serine and instead led to secretion of glycolate. We hypothesized that *cyb2Δ K. phaffii* strains with an intact CBB cycle could be used as RuBisCO oxygenation biosensor strains where the relative levels of glycolate secretion would correlate with the levels of oxygenation of the chosen RuBisCO enzymes. Four different RuBisCO proteins were tested in the autotrophic *CYB2* deletion strain and growth as well as glycolate production was monitored (Fig. 6). The type II RuBisCO proteins of *T. denitrificans* and *Hydrogenovibrio marinus* resulted in fastest growth and similar glycolate production. The RuBisCO protein of *Gallinella sp*. could only barely facilitate growth on CO2 in our synthetic autotrophic strain. In addition, only small amounts of glycolate were produced. Switching to the type I RuBisCO of *C. necator*, which was tested together with the co-expression of its corresponding RuBisCO activase *CBBX* resulted in the lowest glycolate production per biomass (**Fig. 6 B**). *C. necator* RuBisRO led however to a lower growth rate than the best performing RuBisCO variants. Still this type of RuBisCO resulted in the lowest specific glycolate production per biomass.

**Figure 6:**
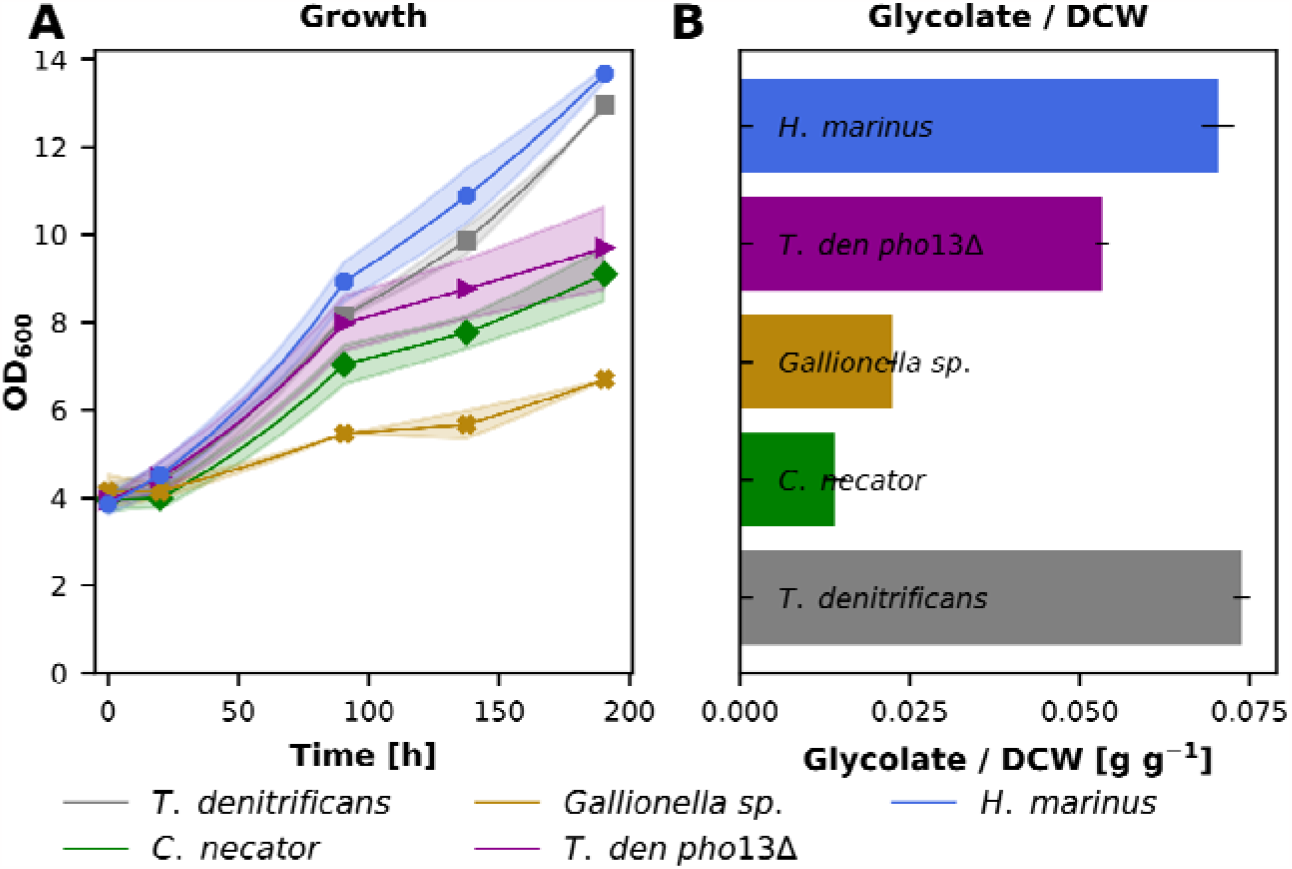
Testing the capabilities of the deletion of *CYB2* as oxygenation biosensor. Autotrophic *K. phaffii* strains using 4 different RuBisCO proteins, and a pho13Δ variant strain of the T. denitrificans RuBisCO strain were tested for (A) growth and (B) the ratio of glycolate to dry cell weight at the last time point. Cultivations were performed at 30°C and 5% CO_2_ in the atmosphere. Solid lines indicate the means, and shades (for A) and error bars (for B) the standard deviation of 3 biological replicates.

Also, a PHO13 knockout strain was included in this series of experiments. This gene was the best BLAST hit of *K. phaffii* genes against *Arabidopsis thaliana PGLP1*, a 2-phosphoglycolate phosphatase. The deletion of this potential phosphoglycolate phosphatase resulted in a reduced growth rate compared to its parental strain. Glycolate production was also reduced, but both were not completely abolished, indicating that more than one phosphatase is acting on 2-phosphoglycolate in *K. phaffii*.

## Discussion

Previously we were able to engineer the methylotrophic yeast *K. phaffii* to use CO_2_ as its sole carbon source by the integration of the CBB cycle. The carbon fixation step in this cycle is catalyzed by the RuBisCO enzyme. This enzyme also tends to perform the oxygenation reaction resulting in a loss of carbon fixation efficiency. In this study we characterized our synthetic autotrophic strains in terms of oxygen tolerance and tried to identify the pathway used for phosphoglycolate regeneration in this yeast strain.

Using different oxygen concentrations in the inlet air clearly showed that the oxygen concentration has a significant influence on the growth rate of the autotrophic yeast strains. Reducing the oxygen levels helped the cells to grow faster, probably because of a reduction in the oxygenation reaction of RuBisCO. The results indicate that at least 5% oxygen in the inlet air are needed for efficient growth. When the oxygen concentrations are lower, not enough oxygen is available for sufficient methanol oxidation providing energy for the cells, resulting in reduced growth rates as seen for the cultivations at 2.5% oxygen (**Fig. 2**). Comparing the growth rates and dissolved oxygen concentrations of the fermentations, we can observe that the optimal dissolved oxygen concentration is around 15% (**Fig. 2 C** and **Supplementary Fig. 1C**). In the faster growing engineered version of this strain the optimal oxygen concentration is increased, probably because of the faster accumulation of biomass and therefore higher energy demand of the cells. Reducing the oxygen concentration did not only help to increase growth in our case but is also reported in microalgae (Kazbar et al., 2019). In contrast most plants do not grow better under reduced oxygen conditions (Tisserat et al., 2002).

To get a deeper understanding of the pathways used to recycle the 2-phosphoglycolate produced via the oxygenation reaction of RuBisCO, a ^13^C tracer experiment using fully labeled glycolate was performed. The metabolites which showed early incorporation of ^13^C atoms were glycine and serine. This indicates that most of the phosphoglycolate is recycled and not fully oxidized to CO_2_. The further flux of the label is probably via deamination of serine to pyruvate. Since pyruvate is part of the central carbon metabolism, it is a branch point where the label enters the TCA cycle. To a similar extent the label was found in the glycolytic metabolites PEP, 2-PG, 3-PG and glycerate. Especially by evaluation of the results of the *CYB2* overexpression strain, we can conclude that the route of the pathway is probably via pyruvate – PEP – 2-PG and 3-PG, the latter reentering the CBB cycle. This strain showed a significantly higher incorporation of label into 2-PG compared to 3-PG and glycerate which supports the proposed route and dismisses the route present in plants via serine-hydroxypyruvate and glycerate. The fractions of label of all evaluated metabolites indicate that the first part of this pathway until serine is very efficient, resulting in nearly fully labeled glycine after 3 h in the overexpression strain. The steps of the pathway following glycine and serine seems to be less efficient, and the incorporated label is not only reentering the CBB cycle but also a significant portion of the labeled carbon atoms is being found in the TCA cycle. The labeling data also confirm that *CYB2* is the major gene responsible for the oxidation of glycolate to glyoxylate since only minor incorporation of ^13^C atoms was found in the *CYB2* deletion strain. Testing the glycolate consumption of strains overexpressing *DLD1*, another gene similar to glycolate oxidases, did not result in any glycolate usage and further proves the importance of *CYB2* in this pathway. As in plants, the phosphoglycolate recycling pathway is a multi-compartment pathway starting from the CBB cycle present in the peroxisomes followed by dephosphorylation probably in the cytosol and oxidation of glycolate by *CYB2* in the mitochondria. By evaluation of the GC-TOFMS labeling data of two additional MS fragments of serine we could deduce where the carbon forming serine from glycine comes from. This is best resolved at t=3 h, when about 40% each of the serine is either unlabeled (M0) or labeled with two ^13^C atoms (M2) (Figure 4). In the decarboxylated (DC) fragment of serine only the fraction containing one ^13^C is dominant, indicating that the carboxyl group that was removed derived from glycolate. The labeling pattern of the amino acid backbone fragment (BB) originating from glycine was not changed significantly. These data allow to conclude that methylene-THF, which forms serine from glycine, probably originates from non-labeled precursors. One possible precursor is methanol, which also partly enters the tetrahydrofolate pathway (Mitic et al., 2023, 2022). Later in the cultivation the fraction of M2 of the BB serine fragment increases indicating that labeled glycine is used for methylene-THF synthesis.

Similar to plants the recycling of 2-phosphoglycolate in the synthetic autotroph *K. phaffii* releases 0.5 mol of previously captured carbon in the form of CO_2_ and additionally 0.5 mol ammonia per mol 2-phosphoglycolate. Both lead to a loss of invested energy. In addition, at least one molecule of ATP is needed during the recycling process in the step from pyruvate to PEP. However, the route via pyruvate and PEP is slightly more energy efficient compared to the hydroxypyruvate route since only one ATP contrary to one NADH has to be invested. In plants, the introduction of the glycerate pathway (Kebeish et al., 2007) or the malate cycle (Maier et al., 2012; South et al., 2019) into chloroplasts could improve the crop yields, making these pathways interesting engineering targets to improve the growth rate of the synthetic autotroph *K. phaffii*. In addition, different synthetic pathways have been designed without carbon and nitrogen loss. These pathways still have to be implemented into plants (Ort et al., 2015; Trudeau et al., 2018) but have already been evaluated in algae (Shih et al., 2014).

The labeling data further showed that overexpression of *CYB2* increased the flux of the whole phosphglycolate salvage pathway. Therefore, we tested the *CYB2* overexpression strain also for growth under autotrophic conditions to evaluate if it can increase the growth rate. The results indicate that native *CYB2* expression is already sufficient to enable a phosphoglycolate salvage pathway rate allowing efficient growth using CO_2_ as carbon source since no improved growth was observed with the overexpression of *CYB2*. As the cells do not secrete glycolate to the supernatant during cultivation it can also be assumed that phosphoglycolate recycling is already sufficient in the parental strain.

The deletion of *CYB2* blocked the phosphoglycolate recycling pathway almost completely and led to the secretion of glycolate. This makes the *cyb2Δ* strain a potential candidate as an indicator for the oxygenation rate of different RuBisCO proteins. To prove this hypthesis, different RuBisCO proteins were tested for the correlation of glycolate production to growth in the *CYB2* deletion strain. As expected, the choice of RuBisCO is a key factor for efficient autotrophic growth in the synthetic autotroph *K. phaffii*. In general, the type II RuBisCO proteins led to faster growth compared to the type I protein tested. Only the type II protein from a *Gallionella sp*., for which the highest turnover number so far was reported (Davidi et al., 2020), was not able to facilitate growth on CO_2_ in our synthetic autotrophic strain. Since the strain using this RuBisCO protein barely grows, the resulting glycolate production rates have to be taken with a grain of salt. The oxygenation test platform clearly revealed the difference of specificity between type I and II RuBisCO proteins. Both type II proteins facilitating efficient growth on CO_2_ resulted in an approximately 5 times higher specific glycolate production rate compared to the type I protein tested. This fits well to specificity data measured using *in vitro* assays (Table 1). We could also prove that *PHO13* is involved in the dephosphorylation of 2-phosphoglycolate since its deletion reduced the specific glycolate production rate.

**Table 1:**
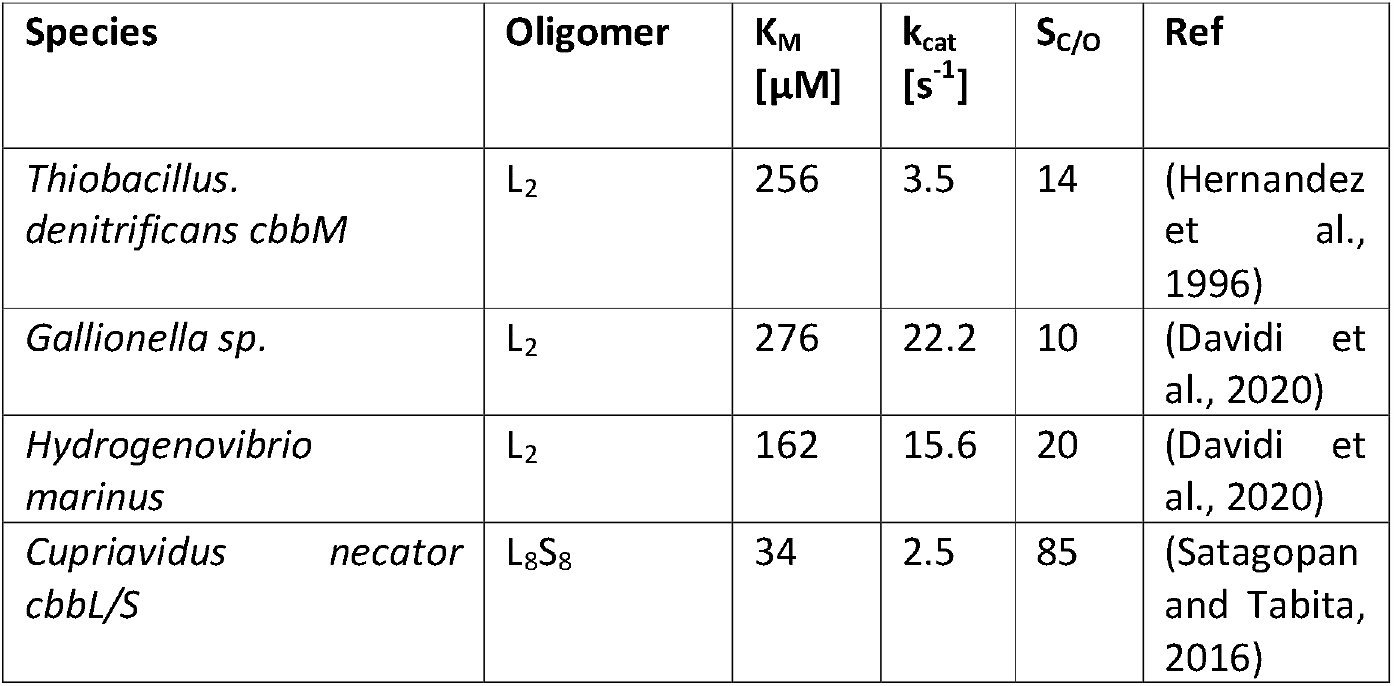
Overview of kinetic parameters of the RuBisCO proteins used in this study (measured using in vitro assays).

| Species | Oligomer | K<sub>M</sub><br>[μM] | k<sub>cat</sub><br>[s<sup>-1</sup>] | S<sub>C/O</sub> | Ref |
| --- | --- | --- | --- | --- | --- |
| *Thiobacillus denitrificans cbbM* | L<sub>2</sub> | 256 | 3.5 | 14 | (Hernandez et al., 1996) |
| *Gallionella sp.* | L<sub>2</sub> | 276 | 22.2 | 10 | (Davidi et al., 2020) |
| *Hydrogenovibrio marinus* | L<sub>2</sub> | 162 | 15.6 | 20 | (Davidi et al., 2020) |
| *Cupriavidus necator cbbL/S* | L<sub>8</sub>S<sub>8</sub> | 34 | 2.5 | 85 | (Satagopan and Tabita, 2016) |

With this work we could prove that balancing the available amount of oxygen can help to improve growth of the synthetic autotrophic *K. phaffii*. We were also able to solve the open question of how synthetic autotrophs can deal with the by-product of the oxygenation reaction and show how versatile a cell’s metabolism can be in responding to newly formed substances and their toxic effects, such as 2-phosphoglycolate from the RuBisCO side reaction. As a next step, other more efficient glycolate salvage pathways like the glycerate pathway could be integrated into our strains to evaluate if they can further improve autotrophic growth rates.

## Material and Methods

### Strain generation (strain list)

The host strain used in this study was the synthetic autotroph *K. phaffii* strain published by Gassler et al. (Gassler et al., 2020). The assembly of all DNA repair templates and over-expression cassettes were done using the Golden *Pi*CS cloning system (Prielhofer et al., 2017) and transformed using the CRISPR-Cas9 system (Gassler et al., 2019). Switching of the RuBisCO protein was done by deleting first the RuBisCO protein of the parental strain followed by the integration of the new RuBisCO protein. Successfully transformed clones were verified by colony PCR. All strains used in this work are given in **Table 2**.

**Table 2:** Overview of the strains used in this study.

| Name | Genotype | Ref. |
| --- | --- | --- |
| Parental strain | CBS7435 *aox1Δ::pAOX1_TDH3*<br>+ *pFDH1_PRK* + *pALD4_PGK1*<br>*das1Δ::pDAS1_RuBisCO* + *pPDC1_groEL* +<br>*pRPP1B_groES das2Δ::pDAS2_TLK1*<br>+ *pRPS2_TPI1* | (Gassler et al., 2020) |
| Eng strain | Parent (PRK 5 C > G) | (Gassler et al., 2022) |
| *cyb2Δ* | Parent + *cyb2Δ* |  |
| *CYB2<sub>medium</sub>* | Parent + *pMDH3 CYB2* |  |
| *CYB2<sub>weak</sub>* | Parent + *pPDC1 CYB2* |  |
| *T. den pho13Δ* | Parent + *cyb2Δ* + *pho13Δ* |  |
| *Gallionella sp.* | Parent RuBisCO (*T. denitrificans*)::<br>RuBisCO (*Gallionella sp.*) |  |
|  | Parent RuBisCO (*T. denitrificans*)::<br>RuBisCO (*H. marinus*) |  |
|  | Parent RuBisCOΔ + *pDAS1 CbbL* + *pFDH1*<br>*CbbS* + *pALD4 CbbX* |  |
| *CYB2(cyt)<sub>medium</sub>* | Parent + *pMDH3 CYB2(del1_117)* |  |
| *CYB2(cyt)<sub>weak</sub>* | Parent + *pPDC1 CYB2(del1_117)* |  |
| *cyb2Δ DLD1<sub>medium</sub>* | *cyb2Δ* + *pMDH3 DLD1* |  |
| *cyb2Δ DLD1<sub>weak</sub>* | *cyb2Δ* + *pLAT1 DLD1* |  |

### Shake flask cultivations

All strains were tested using shake flask cultures. As a first step a preculture on YPG (yeast extract 10 g L^−1^, soy peptone 20 g L^−1^, glycerol 20 g L^−1^) was performed overnight at 30°C and 180 rpm. Afterwards optical density was measured at 600 nm and the number of cells needed to inoculate the main culture (OD of 1, 4 or 20, respectively) was harvested, washed twice with water and resuspended in phosphate buffered Yeast Nitrogen Base Without Amino Acids (YNB) (3.4 g L^−1^, pH 6, 10 g L^−1^ (NH_4_)_2_SO_4_ as nitrogen source and 0.5% (vol/vol) methanol at the beginning as energy source). For the experiments using glycolate as carbon source additionally 0.5 g L^-1^ (for the pre-tests) or 1 g L^-1^ glycolic acid were added. All experiments using glycolate as carbon source were incubated at 30°C, 180 rpm and ambient CO_2_ concentrations. All experiments where cells were grown autotrophically were incubated at 30°C, 180 rpm and 5% CO_2_ in the atmosphere. For all experiments, methanol concentrations were measured on the next day and adjusted to 1%. Afterwards samples were taken regularly and optical density, methanol and glycolic acid concentrations were measured. Water was added to correct the culture volume for evaporation.

### HPLC measurements

HPLC measurements to determine methanol and glycolic acid concentrations, were done according to an already published workflow (Baumschabl et al., 2022).

### Dry cell weight determination

Glass vials were incubated at 105°C for at least 24 hours, cooled down in the desiccator and weighed. Ten mL of culture were harvested by centrifugation, washed twice with water, transferred into the pre-weighed glass vials and incubated at 105°C until the cells were fully dried. After cooling down in the desiccator, the glass vials including the dried cells were weighed again and the dry cell weight was calculated.

### Labeling experiments

To determine the route of the phosphoglycolate salvage pathway, ^13^C labeling experiments were performed. The protocol was similar to the other shake flask cultivations using a starting OD of 20 but fully ^13^C labeled glycolic acid was used (Merck product no. 604011) for all ^13^C cultures and unlabeled glycolic acid with a natural isotopologe distribution for all ^nat^C cultures. Samples were taken at 2, 24 and 48 hours: 500 μL for the determination of optical density, the remaining glycolic acid as well as methanol concentration, and 3 mL for metabolic sampling & GC-TOFMS isotopologue analysis.

### Metabolic sampling & GC-TOFMS isotopologue analysis

The metabolomics workflow was performed as described by Mitic et al. (Mitic et al., 2022) based on the method of Mairinger et al., 2015.

In brief, the cell suspension was rapidly quenched in a 4-fold volume of 60% methanol, 125 mmol L^-1^ TRIS-HCl, 55 mmol L^-1^ NaCl at pH 8.2 and -27°C (Mattanovich et al., 2017) and filtered through a cellulose acetate filter after vortexing the mixture for 4 s. After washing the cells on the filter with 60% methanol, the filters were stored at -70°C. The consecutive boiling ethanol extraction was performed with 4 mL 75% ethanol at 85°C. After centrifugation the supernatants with the extracted metabolites were evaporated until complete dryness with a vacuum centrifuge before reconstitution in 1 mL H_2_O.

For the GC-TOFMS measurements, two methods were applied to cover all needed metabolites in the linear range of the instrument. Automated just-in-time derivatization prior to sample injection was employed to stabilize metabolites and reduce their boiling point. Ethoximation followed by trimethylsilylation was combined with splitless injection and chemical ionization for the analysis of the phosphorylated metabolites and sugar compounds as well as other intracellular metabolites of low abundance (Mairinger et al., 2015). *Tert*-butylsillylation was combined with 1:50 split injection and electron ionization for the analysis of organic acids and amino acids, a method which offers some positional information due to specific fragmentation patterns of the amino acids (Zamboni et al., 2009). The methods used for specific metabolites and samples are listed in in Supplementary information Table S1.

For the evaluation of ^13^C incorporation into the metabolites isotopologue distribution analysis was conducted. The extracted ion chromatograms of all carbon isotopologues of a target analyte were integrated (evaluated mass/charge ratios listed in (Mitic et al., 2022)). The peak areas were corrected with the software ICT correction toolbox v.0.04 for the contribution of other heavy isotopes except ^13^C, as well as the contribution of the natural ^13^C abundance of the derivatization agent. The carbon isotopologue fractions were calculated as follows:

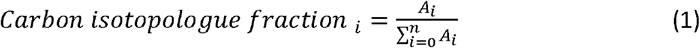

*n* = number of carbon atoms in the metabolite, *A*_i_ = ICT corrected peak area of isotopologue i, *i*.*e*. an isotopologue containing i numbers of ^13^C atoms

For some metabolites such as the amino acids multiple fragments and/or adducts of both measurement methods display the carbon distribution. For data evaluation, the fragment or adduct was chosen based on the trueness of the ^nat^C control sample of the respective sequence (chosen method/ fragment/ adduct listed in Supplementary information Table S1). As the labeling experiments were conducted in biological duplicates, the displayed error bars stem from the standard deviation of duplicates multiplied with the correction factor 1.253314 to compensate for the small sample size (Roesslein et al., 2007). For the ^nat^C control samples average and standard deviation of all data are displayed.

### Bioreactor cultivations

Bioreactor cultivations were performed using 1.0 L DASGIP reactors (Eppendorf). Fermentations were performed using YNB with 10 g L^−1^ (NH_4_)_2_SO_4_ as the nitrogen source, buffered at pH 6 using 100 mmol L^-1^ phosphate buffer, 0.5% (vol/vol) methanol at the beginning as energy source, at 30°C and constant stirrer speed of 300 rpm. Different oxygen concentrations of 5, 10, 15 and 20%, respectively in the inlet gas flow were used to vary the available amount of oxygen, and CO_2_ concentrations were kept constant at 5%.

From an overnight YPG preculture the amount of cells to inoculate 500 mL of culture with an OD of 1 were harvested, washed twice with water and transferred into the bioreactors. The first sample was taken on the next day and methanol concentrations were adjusted to 1% (vol/vol). Afterwards samples were taken daily, and optical density and the methanol concentrations were measured. Every other day the methanol concentrations were adjusted to 1% (vol/vol).

## Author contributions

Ö.A., C.T., S.H., and D.M. designed research; M.B. and B.M.M. performed research; M.B., Ö.A., B.M.M., C.T., and S.H. analyzed data; and M.B., Ö.A., and D.M. wrote the paper.

## Acknowledgments

We want to thank Lisa Lutz for her great help performing some of the experiments. This work was supported by the Federal Ministry for Digital and Economic Affairs, the Federal Ministry for Climate Action, Environment, Energy Mobility, Innovation and Technology, the Styrian Business Promotion Agency SFG, the Standortagentur Tirol, the Government of Lower Austria and ZIT - Technology Agency of the City of Vienna through the COMET Funding Program managed by FFG. We thank the Austrian Science Fund for support to D.M., S.H., M.B. and B.M.M. (FWF W1224, Doctoral Program on Biomolecular Technology of Proteins (BioToP)), as well as support to Ö.A. (FWF M2891). Ö.A. and D.M. were additionally supported by the VIVALDI project which has received funding from the European Union’s Horizon 2020 research and innovation programme under grant agreement No 101000441. This project was supported by EQ-BOKU VIBT GmbH and the BOKU Core Facility Mass Spectrometry.

