## Supplementary File for "A native phosphoglycolate salvage pathway of the synthetic autotrophic yeast *Komagataella phaffii*"

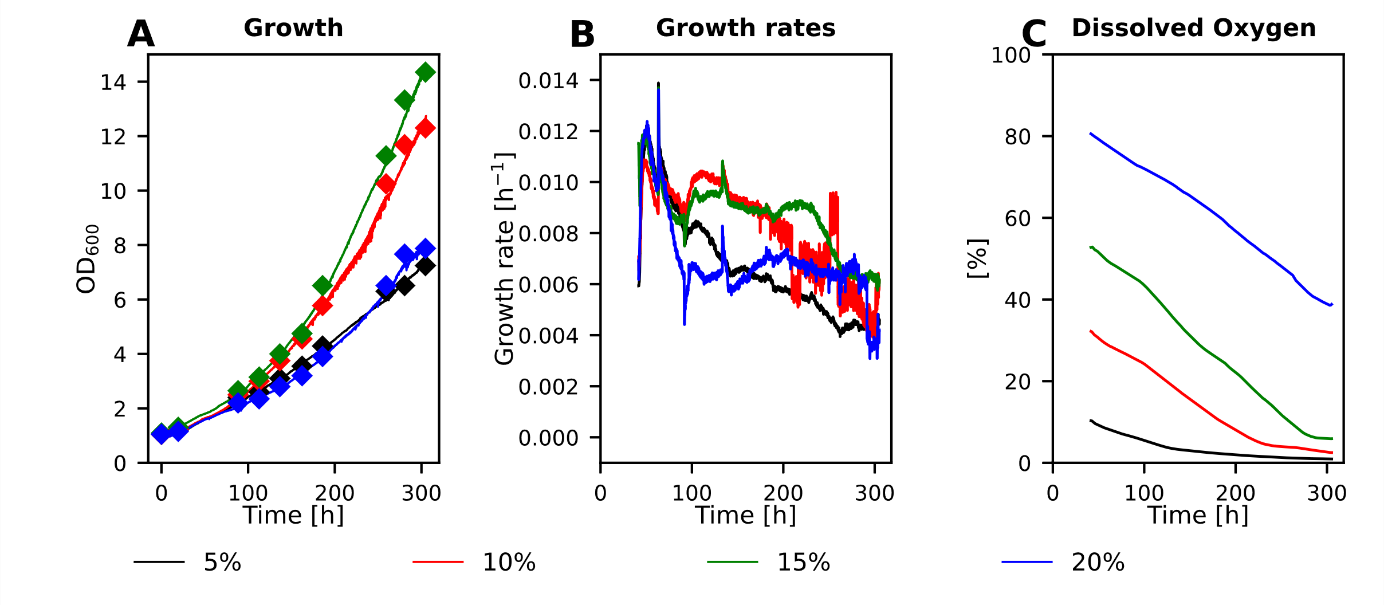

**Supplementary Figure 1:** Cultivations using different oxygen concentrations in the reversed engineered strain. (A) Diamonds: offline OD_600_ measurements and solid line online OD probe to monitor growth, (B) calculated growth rates and (C) dissolved oxygen concentrations of 5 different fermentations using oxygen concentrations in the inlet air from 5% to 20% v/v. Cultivations were performed at 30 °C with a constant stirrer speed of 300 rpm and a CO_2_ concentration in the inlet air of 5 %.

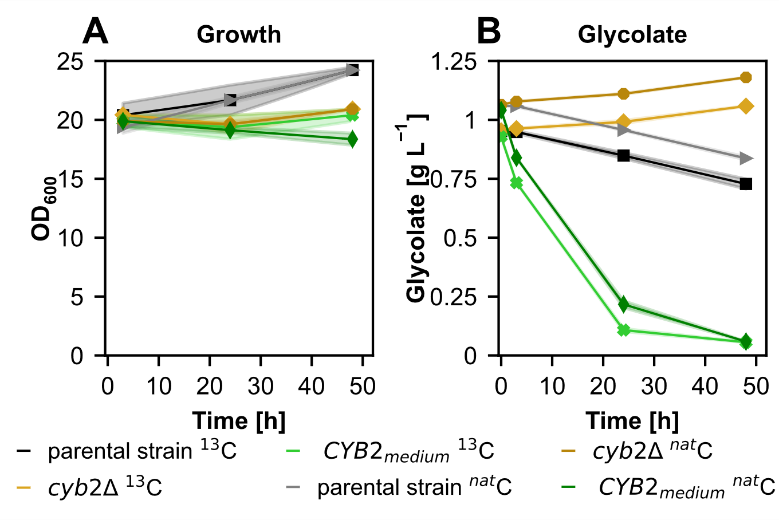

**Supplementary Figure 2:** Growth and glycolate concentrations of the glycolate labeling experiment. (A) Growth and (B) glycolate concentrations in the supernatant of the synthetic autotrophic *K. phaffii* strain (parental strain), a *CYB2* overexpression strain using a medium strength promoter, and the *CYB2* deletion strain cultivated using either glycolate with a natural isotopologue distribution (^nat^C) or fully ^13^C labeled glycolate. Cultivations were performed at 30°C and ambient CO_2_ concentrations in the atmosphere. Solid lines indicate the mean and shades the standard deviation of 2 biological replicates.

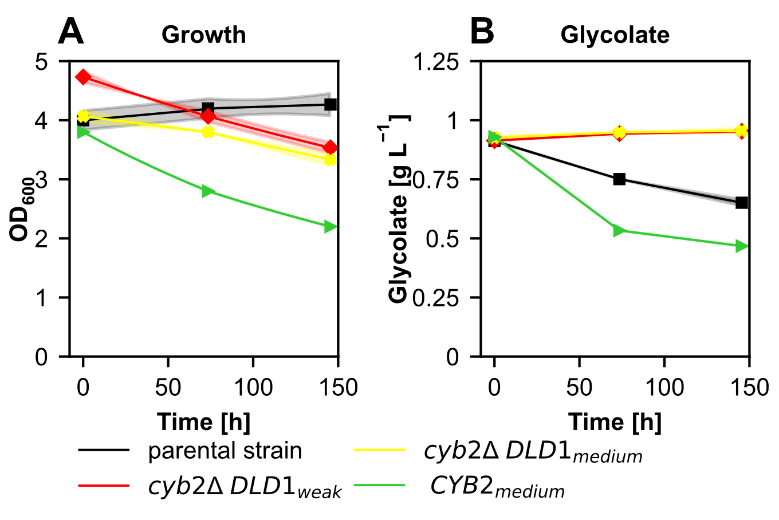

**Supplementary Figure 3:** Can *DLD1* replace the role of *CYB2* in the recycling of 2-phosphoglycolate? (A) Growth and (B) glycolate concentrations in the supernatant of the synthetic autotrophic *K. phaffii* strain (parental strain) and *CYB2* overexpression strains using a medium strength promoter and *CYB2* deletion strains overexpressing *DLD1* using a medium or weak promoter. Cultivations were performed at 30 °C and ambient CO_2_ concentrations in the atmosphere. Solid lines indicate the mean and shades the standard deviation of 2 biological replicates.

**Supplementary Table 1:** List of GC-MS and data evaluation methods used for evaluation of the labeling data.

| **Metabolite** | **GC-MS method** | | **Data evaluation** | | | **Used for** |
| --- | --- | --- | --- | --- | --- | --- |
|  | **Derivatization and Ionization** | **Injection** | **Fragment / Adduct** | **MS data evaluation type** | **Mass extraction window [±ppm]** |  |
| Glycine | TBDMS GC-EI-TOFMS | split 1:50 | [M-CH_3_]^+^ | profile | 50 | Parent T0, CYB OE T0, T1 |
|  | TBDMS GC-EI-TOFMS | split 1:50 | [M-C_4_H_9_]^+^ | profile | 50 | dCYB T1, T2 |
|  |  |  |  | centroid | 50 | Parent T1, T2 |
| Serine | TBDMS GC-EI-TOFMS | split 1:50 | [M-CH_3_]^+^ | centroid | 50 | CYB OE T1 |
|  |  |  | [M-C_4_H_9_]^+^ | profile | 50 | Parent T0, T1, T2, CYB OE T0, dCYB T1, T2 |
| Serine BB | TBDMS GC-EI-TOFMS | split 1:50 | [f302]^+^ | profile | 50 | Parent T0, T1, T2, CYB OE T0, T1, dCYB T1, T2 |
| Serine DC | TBDMS GC-EI-TOFMS | split 1:50 | [M-C_5_OH_9_]^+^ | centroid | 50 | dCYB T1, T2 |
|  |  |  | [M-C_7_O_2_SiH_9_]^+^ | profile | 50 | Parent T0, T1, T2, dCYB T2 |
| PEP | EtOx/TMS GC-CI-TOFMS | splitless | [M+H]^+^ | centroid | 50 | Parent T1, T2 |
| Aspartate | TBDMS GC-EI-TOFMS | split 1:50 | [M-CH_3_]^+^ | profile | 50 | Parent T0, T1, T2, CYB OE T0, T1, dCYB T1 |
|  |  |  |  | centroid | 50 | dCYB T2 |
| Malate | EtOx/TMS GC-CI-TOFMS | splitless | [M+H]^+^ | profile | 50 | Parent T0, T1 CYB OE T1 |
|  |  |  |  | centroid | 50 | Parent T2, CYB OE T0 |
|  | TBDMS GC-EI-TOFMS | split 1:50 | [M-C_5_OH_9_]^+^ | profile | 50 | dCYB T1, T2 |
| 2-PG | EtOx/TMS GC-CI-TOFMS | splitless | [M+H]^+^ | profile | 50 | Parent T2, CYB OE T0 |
|  |  |  |  | centroid | 50 | Parent T1, CYB OE T1 |
| 3-PG | EtOx/TMS GC-CI-TOFMS | splitless | [M+H]^+^ | profile | 50 | Parent T0, T1, T2 |
|  |  |  |  | centroid | 50 | CYB OE T0, T1 |
| Glycerate | TBDMS GC-EI-TOFMS | split 1:50 | [M-C_4_H_9_]^+^ | profile | 50 | Parent T0, T1, T2, CYB OE T0, T1, dCYB T1, T2 |
| R5P | EtOx/TMS GC-CI-TOFMS | splitliess | [M-CH_3_]^+^ | profile | 50 | Parent T1, T2, CYB OE T0, T1 |
| Alanine | TBDMS GC-EI-TOFMS | split 1:50 | [M-C_4_H_9_]^+^ | profile | 50 | Parent T0, T1, T2, CYB OE T0, T1 |
| Fumerate | TBDMS GC-EI-TOFMS | split 1:50 | [M-C_4_H_9_]^+^ | profile | 50 | Parent T0, T1, T2, CYB OE T0, T1, dCYB T1, T2 |
| Glutamate | TBDMS GC-EI-TOFMS | split 1:50 | [M-CH_3_]^+^ | profile | 50 | Parent T1, T2, CYB OE T1, |
|  |  |  |  | centroid | 50 | Parent T0 |
